# Discovering compounds mimicking calorie restriction using mammalian gene expression profiles

**DOI:** 10.1101/2020.06.03.131219

**Authors:** Alexei Vazquez

**Author notes:** Corresponding author, Alexei Vazquez, Cancer Research UK Beatson Institute, Switchback Road, Bearsden, Glasgow, G61 1BD, UK.

## Abstract

Obesity is a risk factor for cardiovascular diseases, diabetes and cancer. In theory the obesity problem could be solve by the adherence to a calorie restricted diet, but that is not generally achieved in practice. An alternative is a pharmacological approach, using compounds that trigger the same metabolic changes associated with calorie restriction. Here I expand in the pharmacological direction by identifying compounds that induce liver gene signature profiles that mimic those induced by calorie restriction. Using gene expression profiles from mice and rat I identify corticosteroids, PPAR agonists and some antibacterial/antifungal as candidate compounds mimicking the response to calorie restriction in the liver gene signatures. Liver gene signature analysis can be used to identify compounds that mimic calorie restriction.

## Introduction

Obesity results from an excess calorie intake relative to expenditure [1]. It seems reasonable that calorie restriction would be sufficient to tackle this problem. However, for a number of reasons, long-term calorie restriction is hard to sustain [2]. A valid alternative is to search for pharmacological agents that could achieve weight loss while maintaining a regular or less restricted diet [3, 4]. Some advances have been made in this direction, with the development of PPARα and PPARγ agonists that stimulate oxidation of fatty acids [5–7].

Drug repurposing is a potential strategy to identify additional compounds to mimic calorie restriction [8]. A popular approach has been to search for compounds that match a specified gene expression profile that has been derived from *in vitro* cell culture systems [9]. Given the proven success of the gene expression profile matching, it is desirable to develop a similar methodology to tackle disease phenotypes that are mainly manifested in mammalian tissues.

Here I propose a gene signature methodology to identify candidate compounds that trigger a target gene signature profile. As a bait I use gene expression profiles from the liver of mice subject to calorie restriction, downloaded from Gene Expression Omnibus. As a probe I use gene expression profiles from the liver of rat exposed to a large collection of compounds from a toxicology study [10]. I identified corticosteroids, PPAR agonists and some antibacterial/antifungal agents as candidate compounds to mimic calorie restriction.

## Materials and Methods

### Gene signatures

All gene signatures were obtained from public repositories or literature reports. The source and gene list of each signature is reported in Supplementary Table 1.

### Gene expression profiles

The gene expression profiles were downloaded from Gene Expression Omnibus. In all cases gene expression was quantified using microarrays and the RMA signal (in log_2_ scale) was used as the gene expression readout.

### Gene set enrichment analysis

The significant induction or repression of a given gene signature on a given sample was quantified using Gene Set Enrichment Analysis (GSEA) [11] as previously described [12]. GSEA results in a positive and a negative score quantifying the induction or repression of the gene signature, together with their associated statistical significance (here 100,000 permutations of the gene assignment to probes). A sample was defined positive (red) for a signature whenever it manifested a significant positive score (statistical significance ≤0.05), negative (blue) whenever it manifested a significant negative score (statistical significance ≤0.05) and no significant change (black) otherwise.

### Similarity statistics

Given the signature scores for two samples, at the red/black/blue level, the similarity was calculated using the Pearson correlation coefficient (PCC). Given two groups of samples associated with two different perturbations the PCC was calculated between all possible pairings between the two groups and the values above the 5%, 50% (median) and 95% of all calculated PCCs were stored for further analysis.

### Positive hits

Given a bait gene signature scores, a probe gene signature score was deemed a positive hit if the 5% value was positive. For each compound tested in our probe dataset, there were several conditions using different doses and treatment durations. Each of these conditions was processed as an independent test. A compound was deemed a hit if it has a significant enrichment of positives across the conditions where it was tested, given the total number of conditions tested and the total number of conditions scoring positive in the whole dataset, using a hypergeometric distribution (Supplementary Table 2).

## Results

### Gene signatures selection

Here I will use liver gene expression profiles as a mean to identify compounds mimicking calorie restriction. These profiles report the expression of more than 20,000 genes as measured using microarrays or RNA sequencing. This large number of variables could carry as a consequence overfitting towards any given dataset and makes the biological interpretation of any result difficult. There are multiple approaches to reduce the number of variables. The standard choice are unsupervised methods such as Principal Component Analysis and its non-linear generalisation using auto-encoders. However, they will be biased for the datasets used to determine the principal components. Analysis of gene signatures provides an alternative supervised approach, with the additional advantage of improving on the biological interpretation.

Here I will adopt a gene signature approach. Gene signatures are gene lists based on pathway annotations, genes changing their expression under pre-defined perturbations, or any biological mean associating a group of genes. Ironically the number of gene signatures in repositories like the Molecular Signatures Database (MSigDB) [13] is getting close to 20,000, bringing us back to the overfitting problem. To tackle this issue, I will limit the analysis to a restricted set of signatures with relevance to calorie restriction (Supplementary Table 1).

The list starts with two fundamental processes of organ homeostasis, cell proliferation and tissue remodelling, followed by gene signatures associated with central metabolism, fatty acid metabolism, cholesterol metabolism and one-carbon metabolism. I will use another subset of signatures to interrogate the potential activation of relevant transcriptional programs. This includes MYC targets as an additional readout for cell proliferation, HIF1α targets for hypoxic response, NFKβ targets for inflammation, ATF4 targets for amino acid stress response, NRF2 targets for oxidative stress response, HSF1 targets for proteotoxic stress response and TFEEB targets for autophagic response. The list of signatures ends with variants of cell death, including apoptosis, necrotic cell death and ferroptosis, plus DNA repair as a readout for DNA damage response.

### Gene signatures validation

The signatures of cell proliferation and tissue remodelling have been tested in the context of human tumours [12]. The metabolic signatures are based on stablished gene annotations of metabolic pathways. The transcription factor targets’ signatures have been developed for the most part from experiments with cells in culture. Therefore, it is important to determine whether they reflect their associated biology in whole tissues. To tackle this problem, I searched the Gene Expression Omnibus database for gene expression profiles associated with the relevant biological conditions.

I identified a dataset reporting liver gene expression profiles of mice in a hypoxic chamber (6-8% oxygen) for two hours (GSE17880, [14]). In this dataset, there is a significant induction of the gene signature of HIF1α targets in the liver of mice under hypoxia relative to controls (Fig. 1A). Interestingly, there is some variability with respect to the metabolic signatures in the group of mice exposed to hypoxia. Yet, in both groups there is a consistent induction of the HIF1α targets signature.

**Figure 1.**
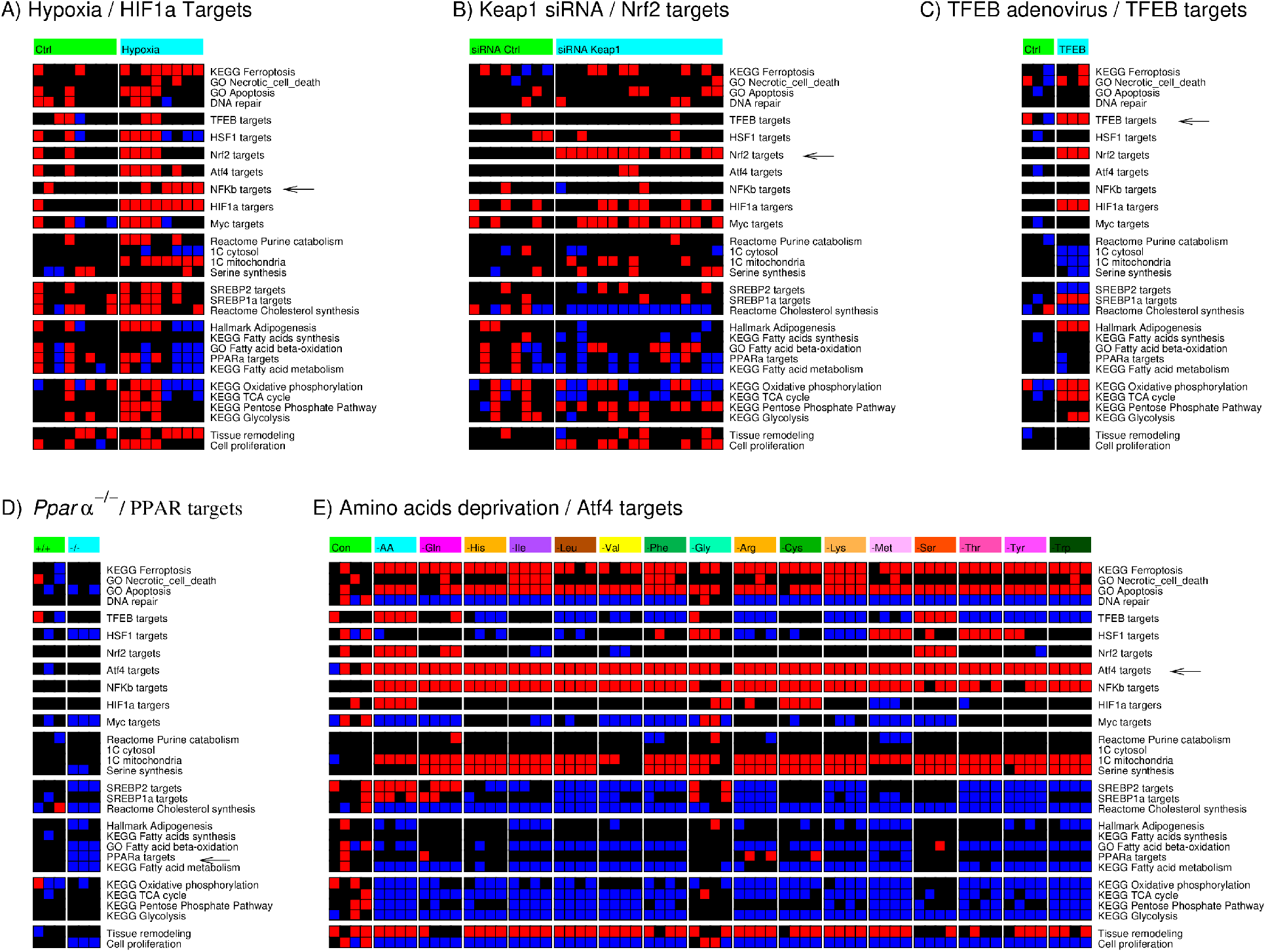
Gene signatures validation. Mouse liver gene signature profiles following pre-defined perturbations. A) Mice in a hypoxic chamber (6-8% oxygen) and controls (21% oxygen). B) Transfection of liver specific siRNA targeting *Keap1* or scrambled control (Ctrl). C) Human *TFEB* expressing adenovirus injection into mice liver and non-injected controls. D) Livers of Wild-type (+/+) and *Pparα^−/−^*(−/−) mice. E) Human MCF7 breast cancer cell lines under amino acid deprivation and control (Ctrl) cells cultured in complete medium. The arrow points to the gene signature that should be activated based on the perturbation applied. Red represents significant induction, black no change and blue significant repression relative to controls (left column).

I identified another dataset reporting liver gene expression profiles of mice injected with scrambled or liver specific siRNA against *Keap1*, the gene encoding the main negative regulator of NRF2 (GSE80956, [15]). I found a clear induction of the NRF2 targets gene signature in the mice treated with the liver specific *Keap1* siRNA (Fig. 1B).

I identified a third dataset where the human *TFEB* gene was injected into the liver of mice using an adenovirus transduction system (GSE35015, [16]). Here again there is the matching induction of the TFEB targets gene signature in mice injected with human *TFEB* expressing adenovirus relative to non-injected controls (Fig. 1C). In this context there is also an induction of the HIF1α and NRF2 targets signatures. This could be biologically relevant. TFEB induces an autophagy program that can lead to mitophagy and the associated reduction in mitochondrial content. The latter could in turn trigger a hypoxic and oxidative stress response, leading to the induction of the HIF1α and NRF2 targets signatures. Whether that is actually the case is beyond the scope of this work.

I also took advantage of the exogenous *TFEB* expression dataset containing liver gene expression profiles of *Pparα^−/−^* mice. In the liver of these mice there is a downregulation of the PPARα targets signature as would be expected (Fig. 1D). In addition, the liver of *Pparα^−/−^* mice exhibit downregulation of the fatty acids β-oxidation gene signature, one of the main processes induced by PPARα.

Unfortunately, I could not find any liver gene expression profile suitable to test the ATF4 targets signatures. In this case I rely on gene expression profiles of the human breast cancer cell line MCF7 cultured under amino acid deprivation (GSE62673, [17]). There is a consistent induction of the ATF4 targets signature under deprivation of each amino acid relative to cells cultured in complete medium (Fig. 1E). Serine synthesis and mitochondrial one-carbon metabolism, two pathways under the control of Atf4 as well [18–20], were also induced upon amino acids deprivation (Fig. 1D).

### Validation across independent experiments

In a second validation I evaluated the consistency of the signature profiles across two independent experiments testing the same perturbation in the same organism. To this end I identified two liver gene expression profiles (GSE21329 [21] and GSE57815 [10]) of rats treated with the PPAR agonists pioglitazone and troglitazone. After computing the significance scores for each gene signature on each liver, I calculated the Pearson correlation coefficient between the gene signature profiles of different experiments for each compound. For pioglitazone I obtained a median correlation of 0.37 with confidence interval [0.16, 0.49] and for troglitazone a median of 0.32 with confidence interval [0.20, 0.44]. The concordance between the signature profiles is also visually observed (Fig. 2A and B).

**Figure 2.**
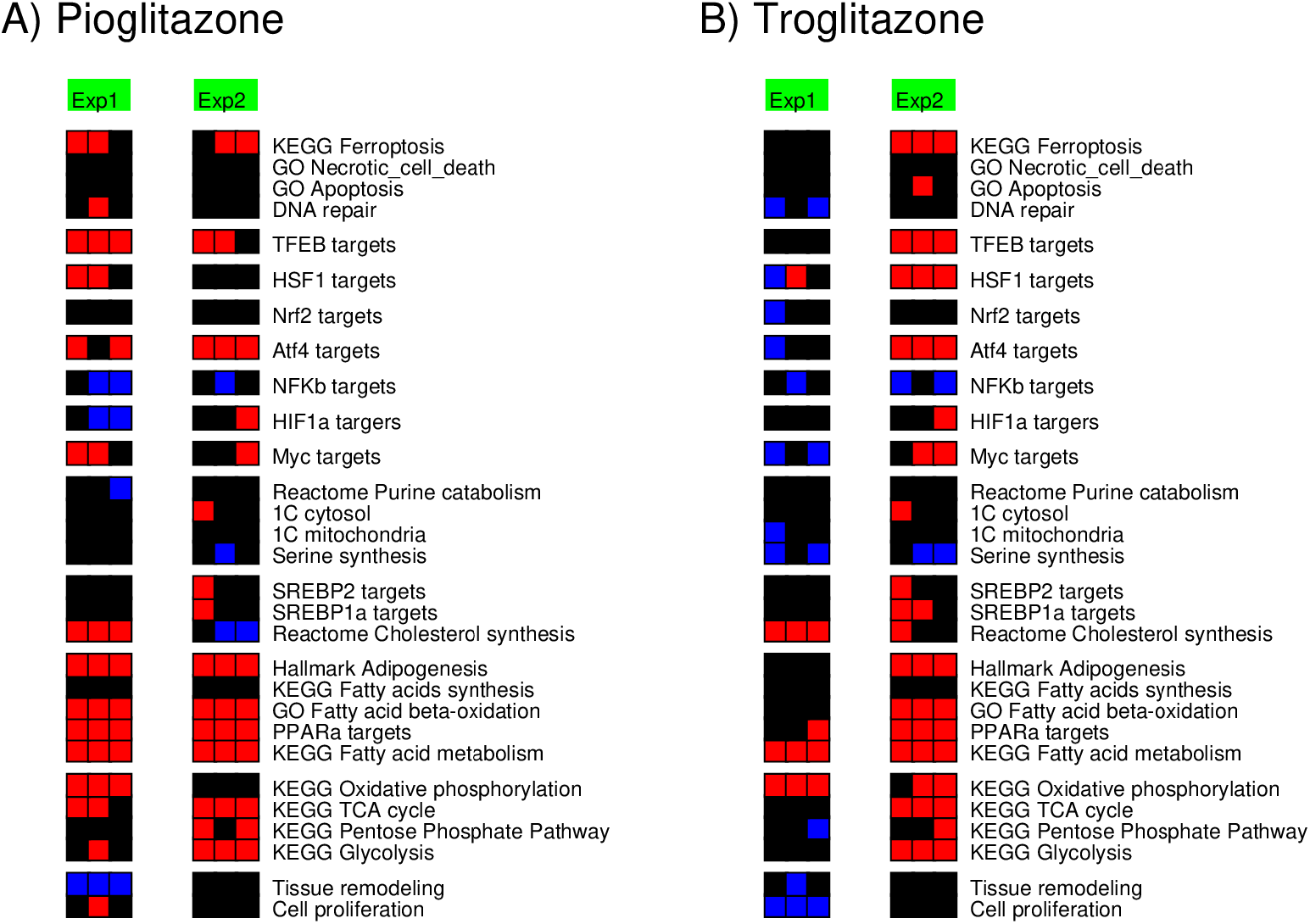
Validation across independent experiments. Liver gene signatures of rats exposed to the PPAR agonists pioglitazone and troglitazone, based on two independent experiments, GSE57817 (Exp1) and GSE21329 (Exp2). Red represents significant induction, black no change and blue significant repression relative to controls (left column).

One could argue that while these correlations have confidence intervals on the positive side they are not too high. However, we should bear in mind that there are differences in the protocols utilised by each independent experiment. One study (GSE59927) used the common rat, higher compound doses (pioglitazone 1,500 mg/kg and troglitazone 1,200 mg/kg) and short treatment time (3 days). The other study (GSE21329) used Zucker obese rats, lower doses (pioglitazone 10 mg/kg and troglitazone 200 mg/kg) and longer treatment time (21 days). Given these protocol differences I argue that median Pearson correlation coefficients in the range of 0.44-0.49 are high.

### Gene signature profile of calorie restriction

Now I switch the attention to the bait, the gene signature profile of calorie restriction. To this end, I selected datasets from Gene Expression Omnibus reporting gene expression profiles of mice subjected to dietary restriction (GSE51885 [22]) or calorie restriction of different magnitude and duration (GSE50789 [23] and GSE40936 [24]).

The gene signature profiles exhibit some similarities across the different experiments and across mouse strains (Fig. 3A). There is a consistent induction of the gene signatures associated with central metabolic pathways (glycolysis, pentose phosphate pathway, TCA cycle and oxidative phosphorylation). However, there is some variability for the gene signatures associated with fatty acid metabolism (fatty acid metabolism, PPARα targets, fatty acid β-oxidation, adipogenesis), cholesterol metabolism (cholesterol metabolism, SREBP1α targets, SREBP2 targets) and one-carbon metabolism (1C cytosol, 1C mitochondria).

**Figure 3.**
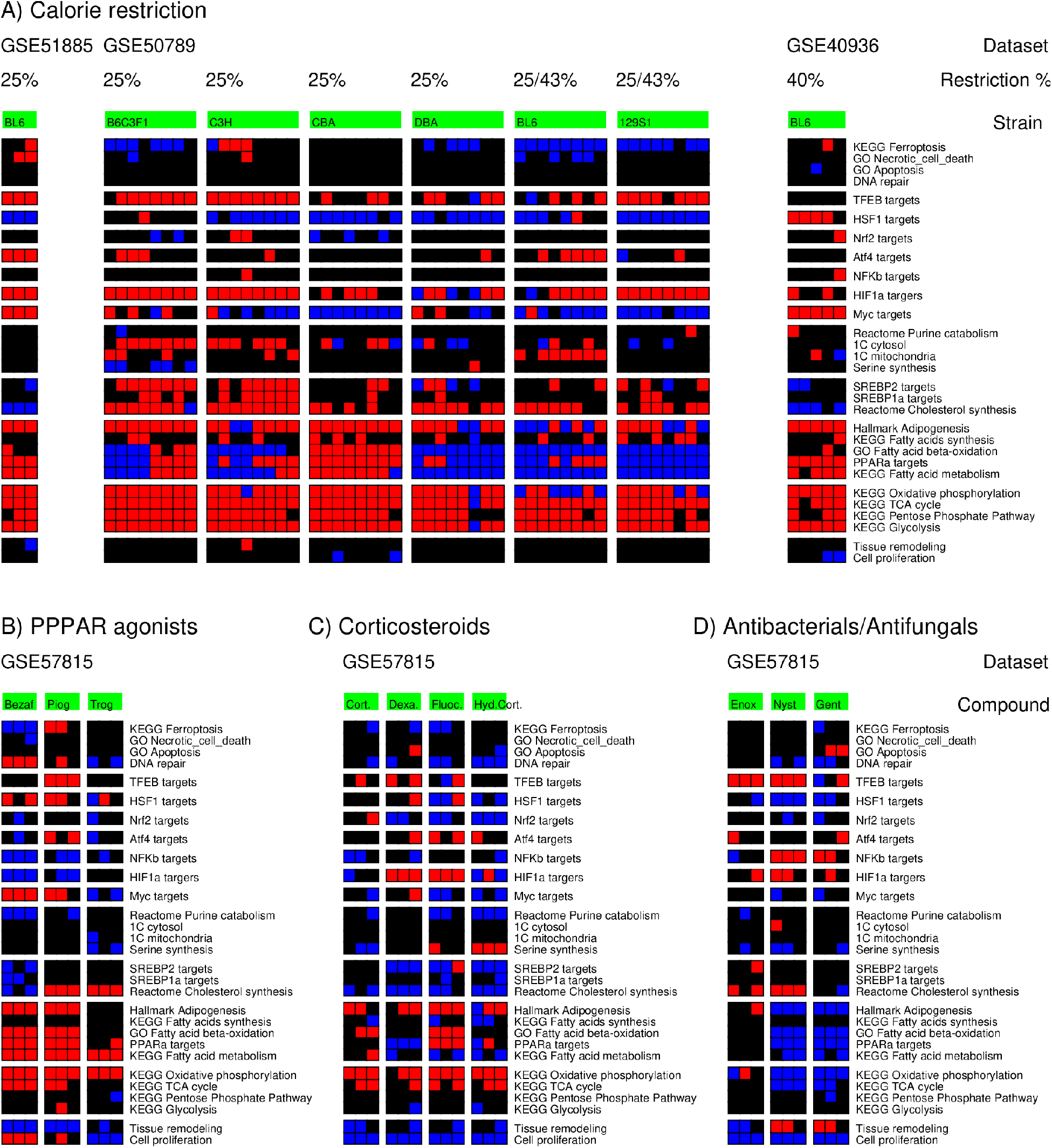
Candidate compounds. A) Liver gene signatures of mice subjected to the reported calorie restrictions. Red represents significant induction, black no change and blue significant repression relative to controls (left column). B-D) Liver gene signatures of mice exposed to PPAR inhibitors (3 days with Bezafibrate 100 mg/kg (Bezaf), Pioglitazone 1,500 mg/kg (Piog), Troglitazone 1,200 mg/kg (Trog)), (B), corticosteroids. (Dexamethasone 1 mg/kg (Dexa), Fluocinolone Acetonide 2.5 mg/kg (Fluoc), Hydrocortisone 56 mg/kg (Hyd Cort)) (C) and antibacterial/antifungal agents (Enoxacin 100 mg/kg (Enox), Nystatin 134 mg/kg (Nyst) and Gentamicin 267 mg/kg (Gent)) (D).

Among the transcription factor signatures, the calorie restriction induces a consistent activation of the HIF1α targets signatures and of TFEB targets signature (except for the GSE40396 dataset). There is a consistent lack of effect on the NFKβ, ATF4 and NRF2 targets signatures. There is a consistent downregulation of the MYC and HSF1 targets signature indicating a suppression (except for the GSE40396 dataset). As noted, the cohort subjected to the more extreme calorie restriction (40%, GSE40396) exhibits a mismatch with respect to the other two cohorts, even when restricting the analysis to the same mouse strain (C57BL/6J).

The GSE51885 dataset, representing dietary restriction with a 25% reduction of all nutrients present in the control diet, exhibits similar gene signatures to the GSE50789 dataset, representing a 25% calorie restriction. This suggests that the calorie restriction component of the dietary restriction is responsible for the observed gene signatures.

Summarising, in the context of mild calorie restriction (~25%), the liver exhibits an induction of gene signatures associated with central metabolism, HIF1α targets and TFEB targets. I note that these changes are consistent with the underlying physiology. It has been shown that autophagy is a modulator of the impact of calorie restriction on longevity [25–27]. The induction of HIF1α targets is most likely a downstream effect, as seen from the injection of *TFEB* expressing adenovirus into the liver of mice (Fig. 1C).

### Compounds matching the calorie restriction gene signature

Having identified the gene signature pattern induced by calorie restriction, we can use it as a bait to uncover candidate compounds to mimic calorie restriction. To this end we need access to liver gene expression profiles following treatment with several compounds. Fortunately, a large-scale toxicology study in rats has profiled 194 compounds (including different vehicles) at different durations (0.25 to 7 days) and reported the corresponding liver gene expression (GSE57815 [10]). I will use the gene expression profiles in this dataset as probes to identify compounds that trigger a gene signature response similar to what observed for calorie restriction.

Table 1 reports the best hits for each dataset and mouse strain. The full list of hits is reported in the Supplementary Table 2. There are three major classes of compounds identified in two or three conditions (dataset/strain): corticosteroids, PPARα/γ agonists and some antibacterial agents. The identification of PPARα/γ agonists is a validation of the methodology. PPARα/γ agonists have been developed for the management of the metabolic syndrome [28]. The identification of corticosteroids and antibacterial/antifungal agents provides two new classes of putative agents for the stimulation of a calorie restriction transcriptional program.

**Table 1.**
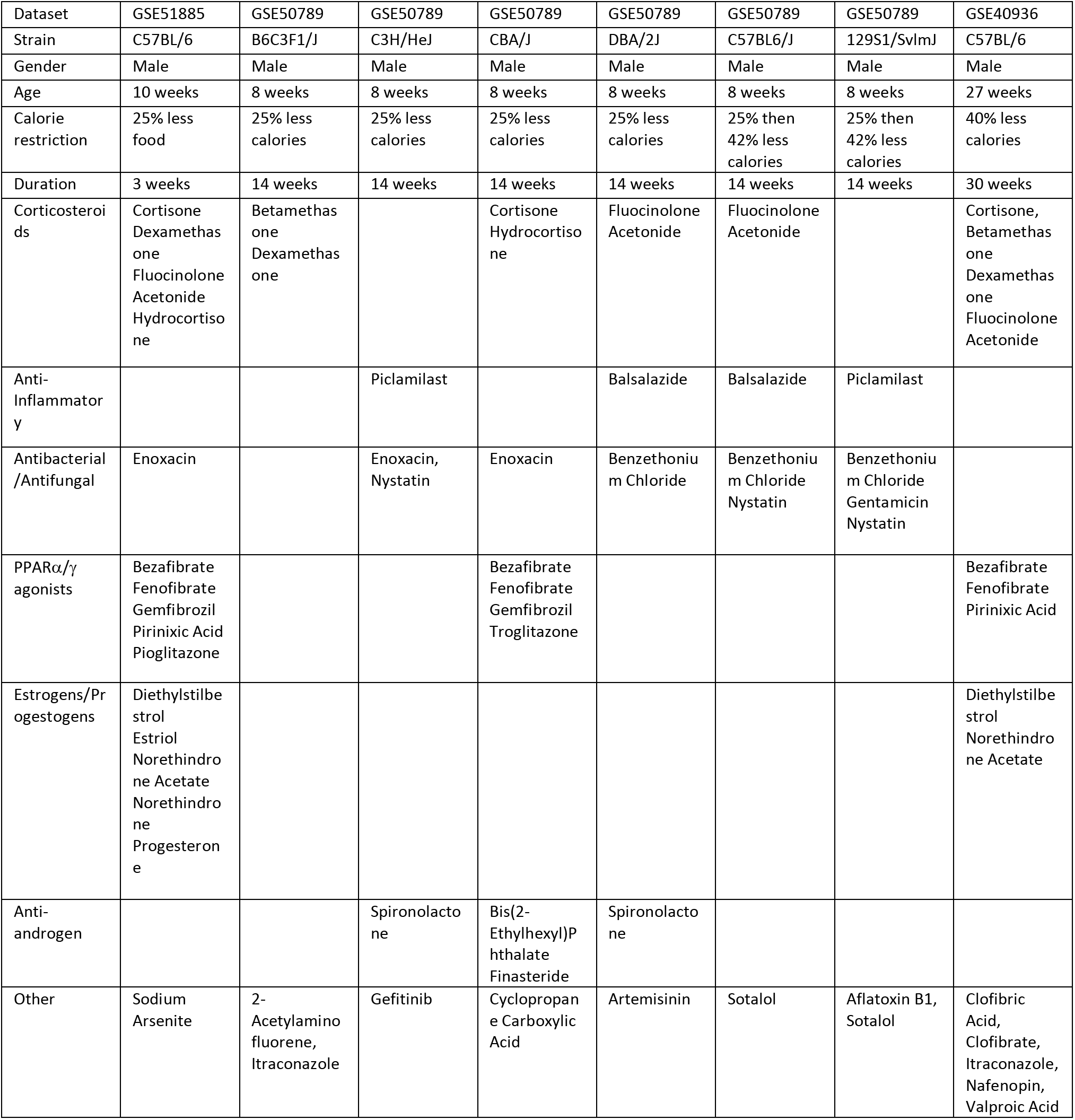
Hit table. Compounds manifesting a positive correlation (95% confidence) across multiple conditions with the calorie restriction gene signature of each bait dataset, grouped by mechanism of action.

The gene signature similarity between calorie restriction and PPARα/γ agonist treatment can be visually inspected in Fig. 3A and B. As calorie restriction, PPARα/γ agonists induce gene signatures of central metabolism (TCA cycle and oxidative phosphorylation), fatty acid metabolism and PPARα targets signatures. However, as a difference with calorie restriction, the PPARα/γ agonists do not induce the gene signatures of glycolysis and the pentose phosphate pathway and only Pioglitazone induces the TFEB targets signature.

Among the compounds scoring the highest I found the synthetic corticosteroids dexamethasone and fluocinolone acetonide, often used as anti-inflammatory agents. The gene signatures associated with these compounds include the induction of HIF1α targets and partially the induction of TFEB targets (Fig. 3C). They also induce some of the signatures of central metabolism (TCA cycle and oxidative phosphorylation) but fail to induce the signatures of glycolysis and the pentose phosphate pathway, as seen for calorie restriction (Fig. 3A).

The other new class of candidate calorie restriction mimics are antibacterial/antifungal agents (Fig. 3D). They induce the gene signatures associated with the TFEB and HIF1α targets, but they tend to induce a downregulation of the gene signature associated with central metabolism. I note that although these compounds are all classified as antibacterial/antifungal agents they have different mechanisms of action. Nystatin is an interesting candidate since it binds to phospholipid membranes and this binding is enhanced by the addition of cholesterol [29]. In fact, nystatin is frequently use as a tool to lower cholesterol and this activity leads to the induction of autophagy [30]. In agreement with these known facts, nystatin induces the gene signature associated with cholesterol synthesis (Fig. 3D).

## Discussion

This work demonstrates that we can use liver gene expression signatures from mammalian model organisms subjected to calorie restrictions as baits to identify compounds that induce a similar transcriptional response. In turn, we can use as probes the liver gene expression signatures induced by treatment with a library of compounds. Here I have used data from the public domain as a proof of concept and identified corticosteroids and some antibacterial/antifungal agents as new candidate compounds to mimic calorie restriction.

In the case of corticosteroids, the list of candidate compounds contains synthetic molecules like dexamethasone and fluocinolone acetonide and endogenous molecules like cortisone. Cortisone is produced from cholesterol in the adrenal gland and then converted to cortisol in the adrenal gland, the liver and other tissues. Cortisone can be supplemented as well. The conversion of cortisone to cortisol in the liver is reduced in obese individuals resulting in lower circulating levels of cortisol [31]. However, the pathophysiology of cortisol in the context of obesity is a complex matter [32], making it difficult to anticipate whether the administration of synthetic corticosteroids would provide any benefit or worsen the disease symptoms. This should be investigated further.

The identified antibacterial/antifungal agents provide additional candidate compounds that should be followed as well. The specific compounds analysed here were constrained to what was included in the toxicology screen. They were not optimised by any means to induce a calorie restriction like response in the liver. The identification of Nystatin, a compound that lowers cholesterol and induces autophagy, highlights a common feature of calorie restriction, the induction of an autophagy gene signature in the liver. Whether that is the most relevant or a leading feature remains to be determined.

There is plenty of room for improvement. One can expand the compounds tested to a library that is more relevant than the one used in the toxicology study. One can fine tune the gene signature list to include other features deemed relevant by human experts or artificial intelligence programs. Finally, the same approach can be deployed to tackle other diseases.

## Acknowledgements

This work was supported by Cancer Research UK A21140 and A17196 (core funding to the CRUK Beatson Institute).

